# Integration of immune cell-target cell conjugate dynamics changes the time scale of immune control of cancer

**DOI:** 10.1101/2024.08.02.606336

**Authors:** Qianci Yang, Arne Traulsen, Philipp Altrock

**Affiliations:** Department of Theoretical Biology, Max Planck Institute for Evolutionary Biology, August-Thienemann-Strasse 2, 24306, Ploen, Germany

## Abstract

The human immune system can recognize, attack, and eliminate cancer cells, but cancers can escape this immune surveillance. The dynamics of these cancer control mechanisms by cells of the adaptive immune system can be captured by variants of ecological predator-prey models. These dynamical systems can describe the interaction of cancer cells and, e.g., effector T cells to form tumor cell-immune cell conjugates, cancer cell killing, immune cell activation, and T cell exhaustion. Target (tumor) cell-T cell conjugation is integral to the adaptive immune system’s cancer control or immunotherapy dynamics. However, it is incompletely understood whether conjugate dynamics should be explicitly included in mathematical models of cancer-immune interactions. Here, we analyze the dynamics of a cancer-effector T cell system regarding the impact of explicitly modeling the conjugate compartment to elucidate the role of cellular conjugate dynamics. We formulate a deterministic modeling framework to compare possible equilibria and their stability, such as tumor extinction, tumor-immune coexistence (tumor control), or tumor escape. We also formulate the stochastic analog of this system to analyze the impact of demographic fluctuations that arise when cell populations are small. We find that explicit consideration of a conjugate compartment can change long-term steady-state, critically change the time to reach an equilibrium, alter the probability of tumor escape, and lead to very different extinction time distributions. Thus, we demonstrate the importance of the conjugate compartment in defining tumor-effector interactions. Accounting for transitionary compartments of cellular interactions may better capture the dynamics of tumor control and progression.

## I. INTRODUCTION

Cancer has been one of the major health and clinical challenges worldwide and is a major contributor to mortality [1]. The human body can command various defenses against cancer growth. Some are cell-intrinsic, such as the apoptotic machinery, which triggers the death of damaged or mutated cells. Tissues can also restrict the growth of developing cancer cell populations, for example, through hierarchical organization into stem, progenitor, and differentiated cell compartments [2–5], or by spatial constraints on the spread of mutants [6, 7]. Apart from these more ‘passive’ cancer suppression mechanisms, the human immune system can recognize, attack, and eliminate cancer cells [8]. During tumorigenesis, cancers may escape from this immune surveillance, but modern therapies can reactivate anti-cancer immune activity [9]. The dynamics of these cancer control mechanisms, e.g., exerted by the adaptive immune system [10], can be captured by predator-prey models and related dynamical systems from ecology [11, 12].

Effector T cells are essential to the human adaptive immune system’s ability to tag and kill aberrant cells [13]. These aberrant tumor cells often generate neoantigens due to tumor-specific molecular alterations, such as mutations, which are recognized as non-self. A neoantigen is a protein appearing on the membrane of immunogenic tumor cells and presents a target to trigger an immune response in the host [14]. Effector T cells can be activated by recognizing neoantigens and killing target cells [15]. This recognition is achieved by the antigen-presenting cells, which display the antigen to provoke the effector T cell’s activation [16], yet also lead to exhaustion in the long run [13]. The activated effector T cell proliferates and uses the T cell receptor on its membrane to recognize and bind to the antigen presented on the surfaces of the targeted tumor cells [17]. The so-formed conjugation enables the effector T cell to deliver a potentially lethal hit to the target tumor cell. Thus, T cell-target cell conjugation is integral to the adaptive immune system’s cancer control or immunotherapy dynamics, as conjugation dynamics may lead to a variation of cytotoxic efficiency [18]. Whether conjugate dynamics should be explicitly captured in mathematical models of cancer-immune interactions is incompletely understood.

Mathematical modeling of biological systems provides powerful tools for formalizing and testing theories [19, 20], explaining empirical observations combining mechanisms that were thought to be unrelated [21–24], and explicitly for understanding and exploring the interaction of the immune system and cancer [25–28]. Kuznetsov et al. introduced a mathematical model of the cytotoxic T lymphocyte response to the growth of an immunogenic tumor [29]. The model involves effector cells, tumor cells, and their formed conjugates that consist of a pair of cells (one tumor or target cell and one effector cell). This conjugate population was then assumed to equilibrate very fast compared to the overall dynamics, subtracting their active contribution to the observed dynamics of tumor burden. Here, we ask under which circumstances the conjugate compartment can assumed to be in such a quasi-steady state in a small tumor. We compare a mathematical model consisting of effector T cells and target tumor cells (two-compartment-model, without conjugate) to a system consisting of effector T cells, target tumor cells, and conjugates (three-compartment-model, with conjugate). We first analyze the deterministic dynamics in these two models: We apply linear stability analysis, investigate the ODE time series damping rate, and quantify the time for the system’s approach to equilibrium. Afterward, we consider stochastic approaches for both models to describe the dynamics in the case of small population sizes. This leads to a formulation of the tumor eradication time and allows us to assess how parameter variability influences the associated time distribution.

## II. METHODS

### A. Deterministic Dynamics

We consider the scheme in Figure 1 (a) to model the interaction of effector T cells and target tumor cells and effector T cell binding with the target tumor cell by matching T cell receptor and proteins on the tumor surface. Subsequently, conjugation either leads to death of tumor cell or the exhaustion of effector T cell.

**Figure. 1:**
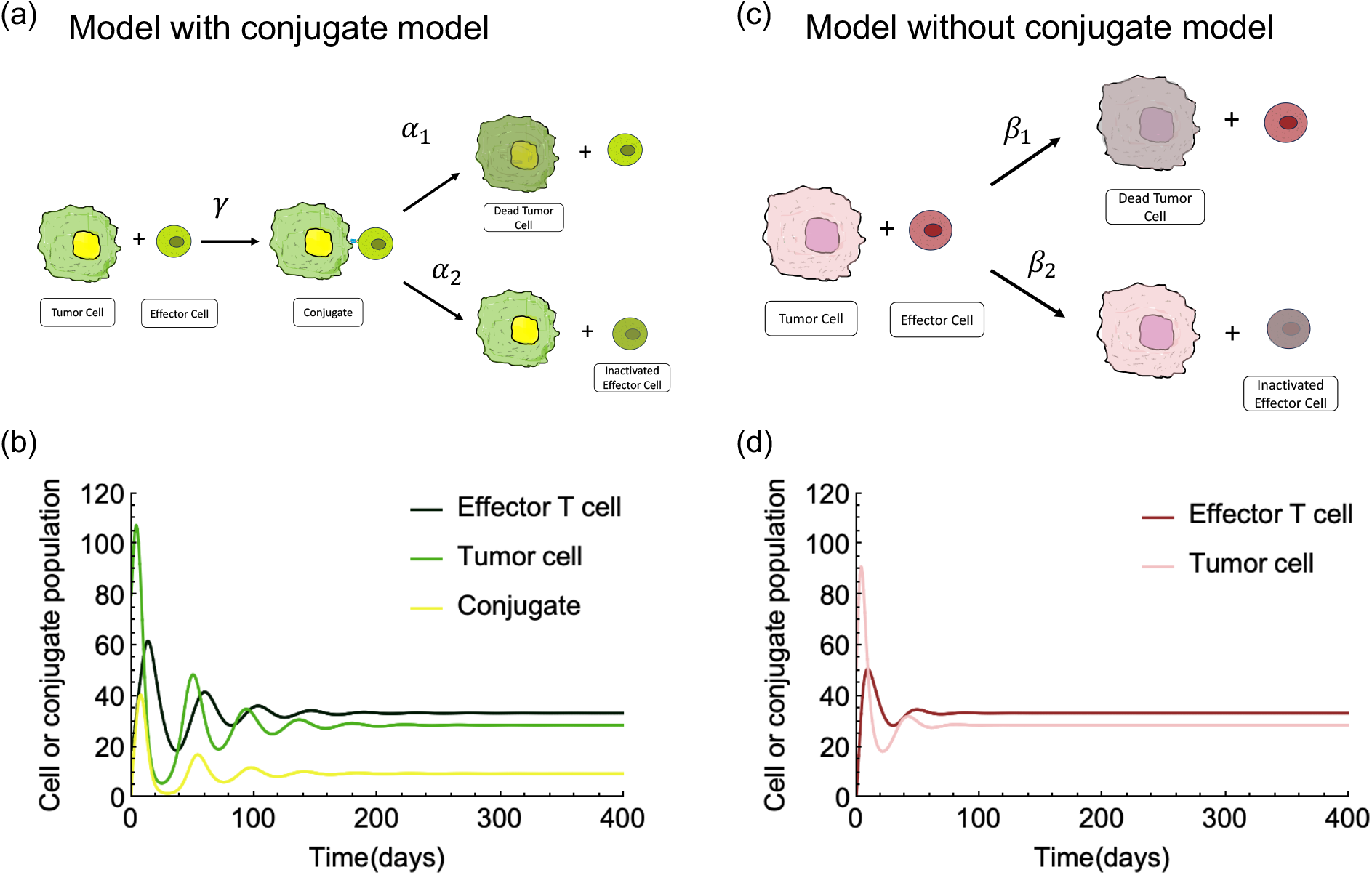
Model with conjugate and Model without conjugate. (a) Schematic drawing of effector T cell, target tumor cell and conjugate system (model with conjugate). (b) Deterministic dynamics of the model with conjugate (parameters *s* = 0.15, *r* = 0.3, *d* = 0.1, *γ* = 0.01, *α*_1_ = 0.9, and *α*_2_ = 0.1). (c) Schematic drawing of the effector T cell and the target tumor cell system (model without conjugate). (d) Deterministic dynamics of the model without conjugate (parameters *s* = 0.15, *r* = 0.3, *d* = 0.1, *β*_1_ = 0.009, *β*_2_ = 0.001).

The model consists of the variables *E* (effector T cell), *T* (target tumor cell), *C* (conjugate), *E∗* (exhausted effector T cell) and *T ∗* (dead tumor cell).

The parameters *γ, α*_1_ and *α*_2_ are non-negative constants: *γ* is the rate of effector T cell and tumor cell conjugate formation, *α*_1_ is the rate at which conjugate leads to tumor cell lysis, and *α*_2_ represents the rate of effector T cell inactivation. Failure to lyse tumor cells results in persistent antigen stimulation. Under such conditions, effector cells gradually become functionally exhausted [30, 31]. The population dynamics of effector T cell and tumor cell is described by

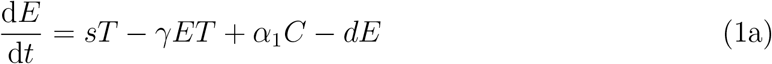

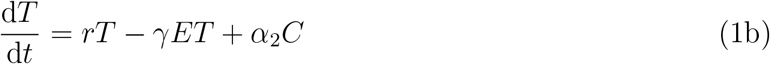

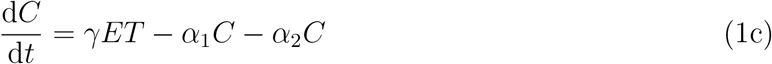

The reproduction term of effector T cells is *sT*. Lanzavecchia and Lezzi [32] observed that naive T cells are stimulated to proliferate under the signaling information conveyed by antigen-presenting cells. This suggests that the influx of mature effector T cells depends on the target tumor cell population *T*, the constant number to naive cells *N* and a constant differentiate rate *σ*. We use a positive coefficient *s* to represent the product of *N* and *σ*, multiplying the population of target cells to express the influx of mature effector T cells. Effector T cells have an elimination term *dE* with positive *d*, due to natural death or migration out of the tumor location. The term *rT* describes the exponential growth of the target tumor cell. The system consists of the effector T cell, target tumor, and conjugate, which is called model with conjugate for short.

Forming a conjugate before target tumor cell lysis and effector T cell exhaustion corresponds to a handling time problem. To investigate whether the handling time affects the dynamics of the system and, if so, to what extent, we modify the model by applying a quasi-steady state approximation, assuming that the dynamics of the conjugate occurs on a much faster time scale compared to those of effector T cell and target tumor cell. This approximation implies setting the differential equation for the conjugate to zero, leading to an expression for the conjugates as a function of E and T

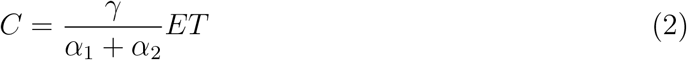

Substituting Eq. (2) into Eq. (1a) and Eq. (1b), we obtain a dynamical system with two compartments (system without conjugate),

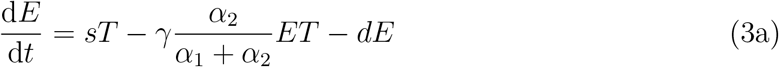

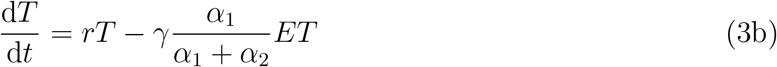

With 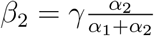 and 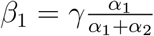, we have a deterministic model for the system without conjugate (see the right part of Figure 1),

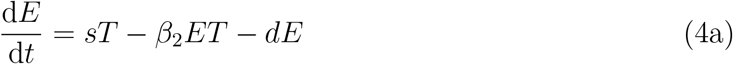

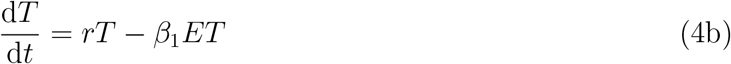

### B. Stochastic Dynamics

The deterministic version of the model describes the dynamic of large cell populations. However, it does not accurately represent the dynamics of small cell populations that can be dominated by stochastic effects. A stochastic model can capture the dynamics for a smaller amount of cells better, as every reaction is explicitly simulated. We implemented the Gillespie algorithm to simulate stochastic equation system trajectories [33]. The Gillespie algorithm is a variant of a dynamic Monte Carlo method and stimulates a realization of the Master equation, which is the first order in time interval of Markov processes in continuous time. Each change of state is the reaction, and the correspondent propensity functions are the likelihood of the reaction happening in time interval. The chemical reactions with corresponding propensity functions for models with conjugate and without conjugate are listed in Table I. The variables *E, T*, and *C* are used for the same populations as in the deterministic model. *E∗* and *T ∗* represent inactivation effector cells and dead tumor cells. In the left part of Table I, parameters *d, s*, and *r* represent the reaction rate of one effector cell’s death, the reaction rate of one effector birth when tumor cells present, and the reaction rate of one tumor cell birth. The parameter *γ* is the reaction rate of one effector cell and one tumor cell binding. The reaction rates that one conjugate leads to either tumor cell being killed or the effector cell inactivation are *α*_1_ and *α*_2_. The propensity functions, also called transition probabilities, are based on the number of cells present in the system. We assume the volume of the system *v* is one. In Table I right table, parameters *d, s* and *r* have the same meaning as those in the left part of Table I. Here, there are two possible events when one effector cell meets one tumor cell, represented by *β*_1_ and *β*_2_. While *β*_1_ describes the reaction rate of effector cell-tumor cell coming together resulting in tumor cell killing, the parameter *β*_2_ is the reaction rate of effector cell and tumor cell coming together resulting in tumor cell survival and effector T cell exhaustion.

**TABLE I:** Table of chemical reactions and corresponding propensity functions for model with conjugate and model without conjugate.

| Chemical Reaction | Propensity Function |
| --- | --- |
| $E \rightarrow \emptyset$ | $dE$ |
| $T \rightarrow T + E$ | $sT$ |
| $T \rightarrow T + T$ | $rT$ |
| $E + T \rightarrow C$ | $\gamma ET/v$ |
| $C \rightarrow T + E^*$ | $\alpha_1 C$ |
| $C \rightarrow T^* + E$ | $\alpha_2 C$ |
| $E \rightarrow \emptyset$ | $dE$ |
| $T \rightarrow T + E$ | $sT$ |
| $T \rightarrow T + T$ | $rT$ |
| $E + T \rightarrow T^* + E$ | $\beta_1 ET/v$ |
| $E + T \rightarrow T + E^*$ | $\beta_2 ET/v$ |

## III. RESULTS

### A. Explicitly modeling conjugate compartments can change long-term steady state

To compare cancer-immune dynamics with and without immune cell-cancer cell conjugation, we first focus on the steady states. Both systems have two steady states, one describing extinction and one coexistence of all described cell types. Note that a condition *α*_1_*s-α*_2_*r >* 0 must be satisfied to make all steady states feasible (i.e. population sizes are not negative). We summarise the stability of steady states in Table II. The trivial fixed points are unstable saddle points in both systems. For the model without conjugate, the coexistence steady state is stable as the eigenvalue of the Jacobian matrix always has negative real part (see Appendix). The stability of the coexistence steady state of the system with conjugate is less straightforward to analyze, as the eigenvalues of the Jacobian matrix are the roots for a cubic polynomial function. Numerical methods are used to determine stability (see Appendix).

**TABLE II:**
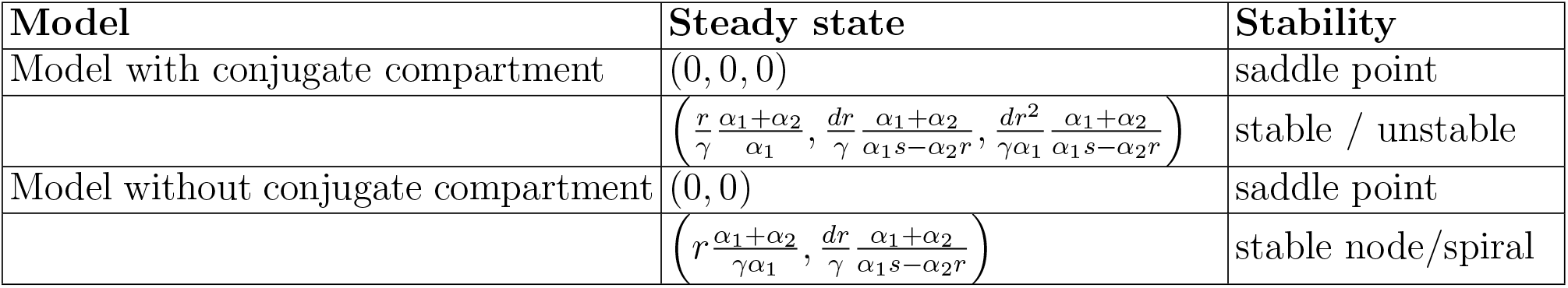
Table of steady states in systems with and without conjugate and corresponding stability behavior.

| Model | Steady state | Stability |
| --- | --- | --- |
| Model with conjugate compartment | $(0, 0, 0)$ | saddle point |
| | $\left( \frac{r}{\gamma} \frac{\alpha_1 + \alpha_2}{\alpha_1}, \frac{dr}{\gamma} \frac{\alpha_1 + \alpha_2}{\alpha_1 s - \alpha_2 r}, \frac{dr^2}{\gamma \alpha_1} \frac{\alpha_1 + \alpha_2}{\alpha_1 s - \alpha_2 r} \right)$ | stable / unstable |
| Model without conjugate compartment | $(0, 0)$ | saddle point |
| | $\left( r \frac{\alpha_1 + \alpha_2}{\gamma \alpha_1}, \frac{dr}{\gamma} \frac{\alpha_1 + \alpha_2}{\alpha_1 s - \alpha_2 r} \right)$ | stable node/spiral |

### B. Taking into account a conjugate compartment affects time scales

From the previous section, the populations for effector and tumor cells have the same steady states for the models with and without a conjugate compartment, but the stability of the coexistence states can differ. In this section, we focus on the time from the initial condition to the two systems’ stable, non-trivial, steady state. We are interested in how the transitional conjugate will influence the time to converge to a stable situation where a small tumor could exist.

In the model with a conjugate compartment, the binding of effector and tumor cells happens first. This process may lead to a longer time to reach a stable coexistence steady state. Two features are important in characterizing the oscillation before the convergence to the fixed point: the amplitude and the damping rate. The oscillation amplitude depends on the difference between initial conditions and the population at steady state. The damping rate depends on the largest real part of the eigenvalues of the ODE’s Jacobian matrix.

The eigenvalues of the Jacobian matrix for the model with a conjugate ODE are too complex to elucidate the relation between them and parameters analytically. A numerical method is more straightforward to reveal which parameter influences the damping rate of the ODE trajectory oscillation. Figure 2 illustrates the relation of the absolute value of damping rate and the parameters: the reproduction rates (*s* and *r*) of the effector cell and the tumor cell, the effector cell death rate *d*, the binding rate *γ*, the tumor-killing rate *α*_1_, and the effector cell inactivation rate *α*_2_. Effector cell and tumor cell binding rate *γ* is the only parameter that does not affect the damping rate. Another interesting parameter is the tumor cell reproduction rate *r*, different from the other parameters, which are positively correlated to the damping rate; the increase of *r* leads to a decrease in the damping rate. This means greater tumor cell reproduction rate results in slower convergence to a co-existence equilibrium. For other parameters, the greater the parameter value, the larger the absolute value of the damping rate - leading to a in faster convergence to a co-existence equilibrium.

**Figure. 2:**
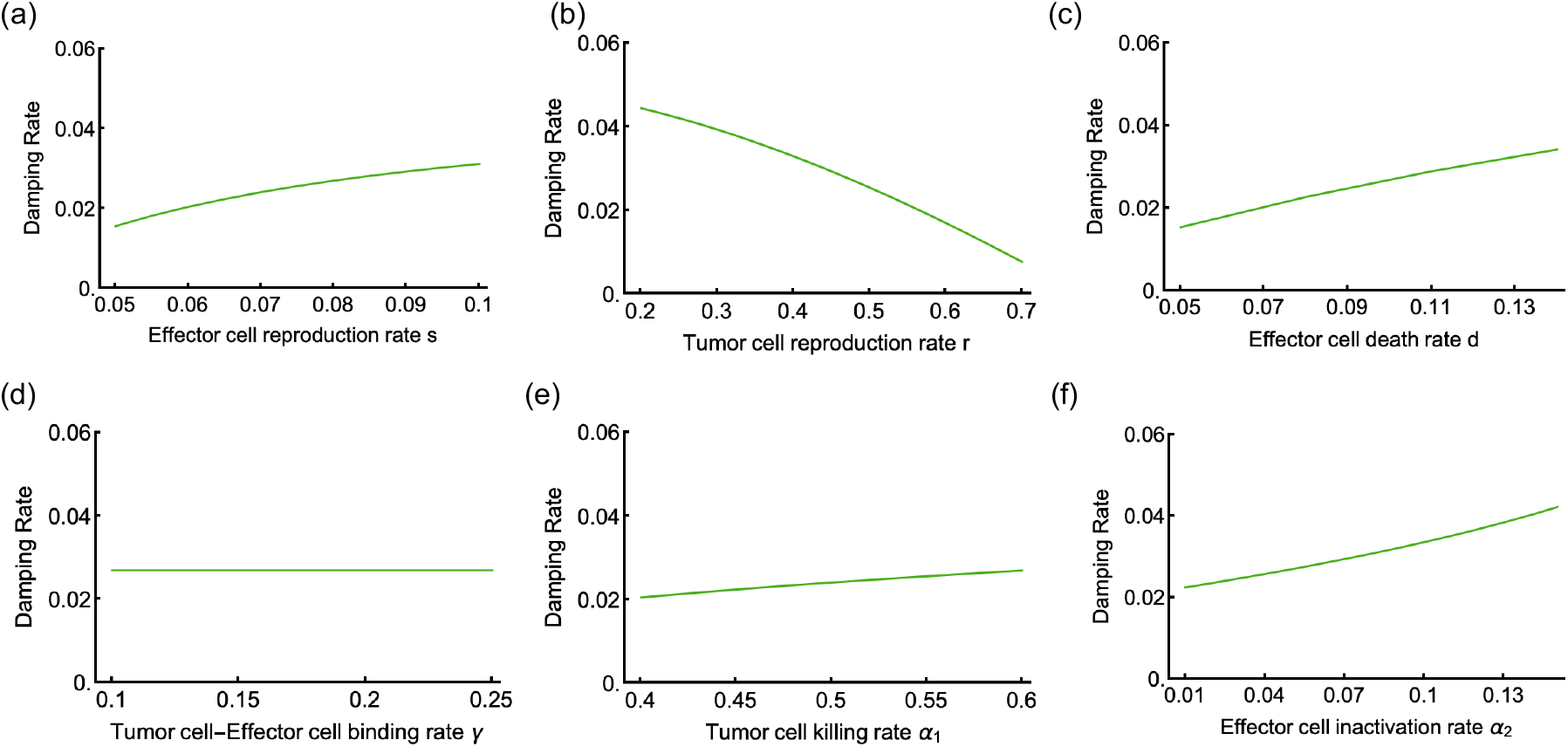
Damping rate in the model with conjugate. The damping rate of the ordinary differential equation trajectories oscillation is the largest real part of the eigenvalues of the Jacobian matrix of the ODE. Six panels show the relation between all six parameters and the absolute value of damping rates. The default six parameters are *s* = 0.08, *r* = 0.15, *d* = 0.1, *γ* = 0.1, *α*_1_ = 0.6, and *α*_2_ = 0.05. In each of the panels, we vary a single one of these parameters: In panel (a) we vary the effector cell reproduction rate *s*, in panel (b) we vary the tumor cell reproduction rate *r*, in panel (c) we vary the effector cell death rate *d*, in panel (d) we vary the effector-tumor cell binding rate *γ*, in panel (e) we vary the tumor cell killing rate *α*_1_, and in panel (f) we vary the effector cell inactivation rate *α*_2_. The damping rate decreases with the tumor cell reproduction rate *r*; independent of the binding rate *γ* and increases with all other parameter changes.

To better visualize the time from the initial condition to the stable coexistence steady state in both systems, we run ordinary differential equations numerical solution under pairs of parameters and record the time until ordinary differential equations trajectories approaches the stable non-trivial steady states. As a criterium to end the numerical solution, we define

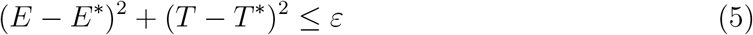

Here that *E*^*∗*^ and *T* ^*∗*^ are the population quantities for effector cell and tumor cell at the coexistence steady state.

We sketch the metric in the phase portrait of the model without conjugate compartment (Figure 3a). When the simulated ODE trajectories come close to the fixed point, the convergence time is the time in which the sum of the squared difference becomes smaller than *ε*. Figure 3b shows the time it takes from the initial condition to the coexistence steady state, choosing different values of 1*/ε*. The convergence time increases with 1*/ε* roughly like a logarithm. This means the time does not increase by the same amount with the reciprocal of *ε* when *ε* is sufficiently small. We choose 0.01 for *ε*. We set the coexistence steady state to be the population at time equals 800 days. Pairs of parameters in the model with conjugate are varied to see their relation to the converge time. Figure 3c presents the convergence time (expressed in color scale) varying tumor cell reproduction rate *r* and effector cell death rate *d*. The convergence time increases with the tumor reproduction rate; this also validates the result that the damping rate is negatively correlated with the tumor reproduction rate, thus leading to slower convergence. Differently, the growth of the death rate of effector cell *d* results in the decrease of converge time. The negative influence on convergence time happens for tumor killing rate *α*_1_ and the effector cell inactivation rate *α*_2_, and they are displayed in Figure 3e. Compared to the model with conjugate, tumor cell reproduction rate *r* plays a different role in converging time in the model without conjugate. With the decline of *r*, it takes a longer time to reach the fixed point. Effector cell death rate *d* plays a consistent role in both systems, as Figure 3d shows. Tumor cell killing rate *β*_1_ is positively correlated with time to converge, while effector inactivation rate *β*_2_ is not. Figure 3 panel (f) depicts the converge time change with variation of *β*_1_ and *β*_2_.

**Figure. 3:**
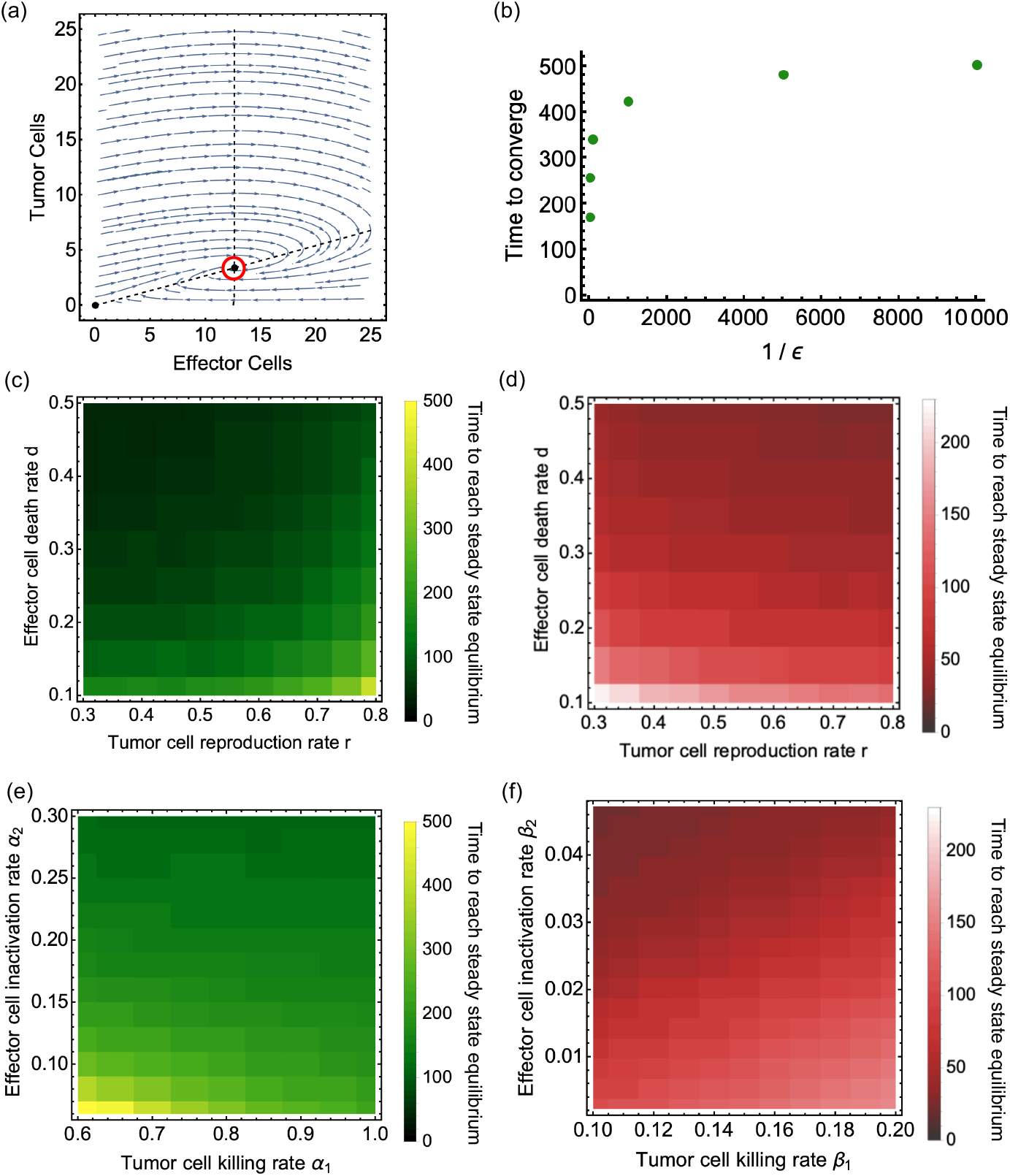
Time to reach the coexistence steady states in both systems. Panel (a) is the 2-dimensional phase portrait of the system without conjugate and a sketch of the threshold *ε* (red circle). When the simulated ODE trajectories approach the threshold, the time it takes from the start to reaching the red circle is recorded as the time to converge. Panel (b) shows the time to converge as a function of 1*/ε*. We take the time points from the initial condition (*E* = 100, *T* = 500) to reach the threshold around non-trivial coexistence steady state in the model with conjugate. Parameters setting: *s* = 0.6, *r* = 0.8, *d* = 0.2, *γ* = 0.08, *α*_1_ = 0.5, *α*_2_ = 0.1. We choose *ε* = 0.01 for the following. Panel (c) and (e) depict the time from the initial condition to reach the threshold around the coexistence steady state in the model with conjugate varying parameters. For (c), *s* = 0.6, *γ* = 0.08, *α*_1_ = 0.8, *α*_2_ = 0.1. For (e), *s* = 0.6, *r* = 0.8, *d* = 0.2, *γ* = 0.08. Panel (d) is the time to the non-trivial steady state when varying target tumor reproduction rate *r* and effector T cell death rate *d* in the model without conjugate, setting *s* = 0.6, *β*_1_ = 0.071, *β*_2_ = 0.009. Panel (f) is the convergence time in different combinations of tumor cell killing rate *β*_1_ and effector cell inactivation rate *β*_2_ in the model without conjugate, setting *s* = 0.6, *r* = 0.8, *d* = 0.2. The initial conditions for panels (c), (d), (e), and (f) are *E* = 100, *T* = 500, and *C* = 0 for panels (c) and (e).

### C. Conjugate changes tumor extinction time and tumor can escape without conjugate

The tumor and effector cells can deterministically coexist in both system settings and at very low levels, leading to possible stochastic extinction. So, the tumor eradication occurs due to stochastic events that drive tumor cells to extinction. We thus ask whether parameter variability or stochastic nature of the underlying process, or both, is responsible for the distribution of the time to cure and if they behave differently in the two models.

Compared to the deterministic version of the model, the stochastic model can capture the dynamics for a smaller number of cells as every change in population is explicitly simulated. We use stochastic simulations to describe the time distribution of tumor eradication using the. Gillespie algorithm. Figure 4 shows 100 simulations of tumour population trajectories in both models. Panel (c) displays long-time fluctuation until tumor extinction. We set trajectories to be slightly transparent, thus denser color represents more trajectories overlap. Most of the trajectories fluctuate and decrease to extinction before 6000 days. In this simulation, the two tumor and effector cell interaction rates *β*_1_ and *β*_2_ are relatively low; their sum is 0.02. This suggests that under this parameter setting, tumor cell population in the system without conjugate can survive at a low amount for a long time before finally extinction.

**Figure. 4:**
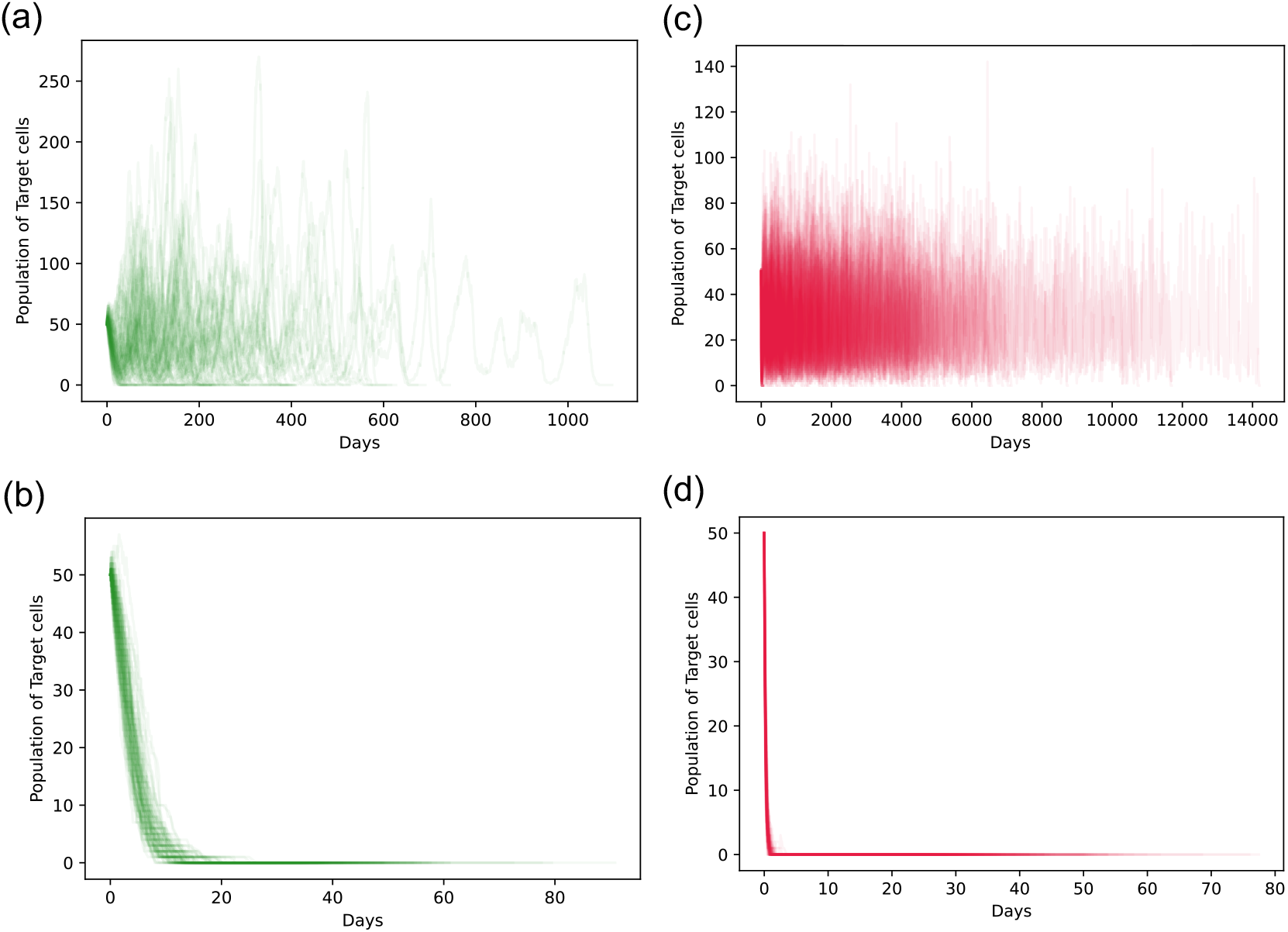
Stochastic simulation results - model with conjugate and model without conjugate tumor trajectories. Panels (a) and (b) describe 100 tumor trajectories in the stochastic simulation results of the model with conjugate. while panels (c) and (d) describe those in the model without conjugate. The initial conditions are effector T cells *E* = 30, target tumor cells *T* = 50. For (a), we choose *γ* = 0.02. In panel (c), we set the parameters *β*_1_ = 0.018 and *β*_2_ = 0.002. In panel (b), *γ* value is increased to 0.2. For panel (d), we set the parameters *β*_1_ = 0.18 and *β*_2_ = 0.02 (the other parameters are *s* = 0.05, *r* = 0.15, *d* = 0.1. In (a) and (b), specially *α*_1_ = 0.4 and *α*_2_ = 0.05).

For the model with conjugate compartment, in most cases the tumor cell population goes to extinction more quickly, as panel (a) in Figure 4 shows. However, the simulation results of both models show a similar trend when a larger value of tumor and effector cell binding rate *γ* is chosen. In panel (b), we choose the same value for all the parameters except increasing tumor and effector cell binding rate *γ* tenfold to be 0.2. In panel (d), the sum of two rates *β*_1_ and *β*_2_ increases to 0.2. An increase of *γ* prevents tumor trajectories from fluctuating; instead, nearly all trajectories decline sharply in the beginning, although trajectories from the model with conjugate reach extinction at a later time. We will investigate how the parameters affect the tumor extinction time distribution.

We compare the tumor eradication time distribution and potential influence parameters in both model settings. We start from the situation where the tumor is in a stable coexistence steady state for both deterministic models and define a criterion to end the simulation naturally. The maximum simulation time is set to be very large, 50000 days. And we either stop the simulation when the maximum time is reached or end it if a stochastic mutation event occurs, at a small rate *µ* = 10^−6^. Six parameters are varied to see if the variability affects the tumor extinction time. Figure 5 show the violin plot of tumor cell eradication time distributions simulated from the model with conjugate. The maximum, minimum, and median are included. We vary the parameters’ value to see the trend and shape of the tumor eradication time distribution. Figure 5 shows that with the increase of effector cell reproduction rate, effector cell - tumor cell binding rate or tumor killing rate will lead to faster the tumor cell population going to extinction, which validates our intuition. The quicker the effector cell is generated, the quicker the tumor cell is bound and killed by the effector cell, and the shorter time the tumor cell population takes to go extinct.

**Figure. 5:**
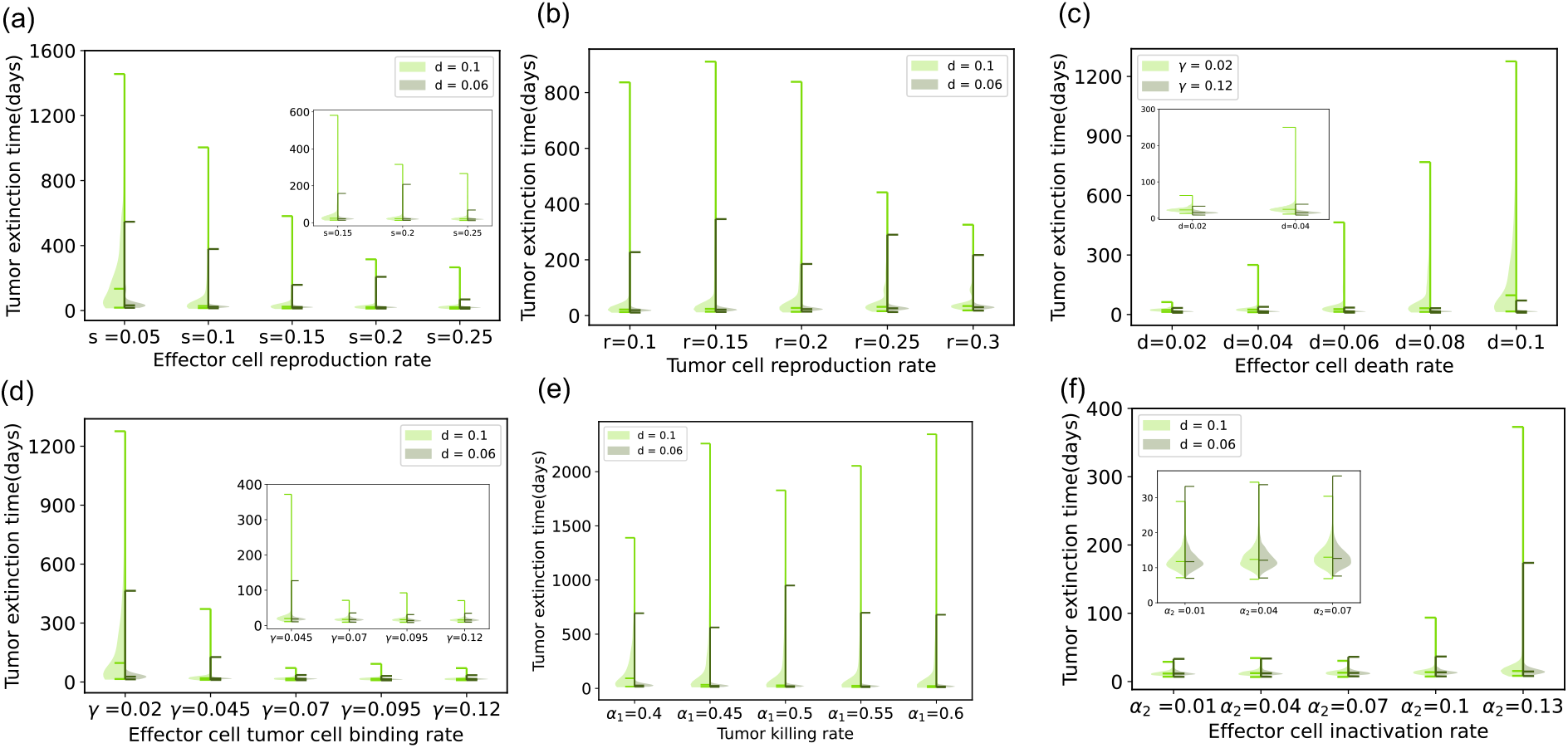
Stochastic simulation results - violin plots of tumor extinction time distribution for model with conjugate. The default parameters are *s* = 0.05, *r* = 0.15, *γ* = 0.02, *α*_1_ = 0.4, and *α*_2_ = 0.05. In the left half part (green color), effector cell death rate parameter is set to be *d* = 0.1 and for the right half (grey color), *d* = 0.06. The initial conditions are effector T cells *E* = 50, target tumor cells *T* = 100. Panel (a) displays tumor extinction time (in days) distribution violin plots for variation of the birth rate of effector cells *s*. In the inset, we zoom into the three distributions on the right. Note that here the tumor killing rate is *α*_1_ = 0.35. Panel (b) displays tumor cell reproduction rate *r* variability and the extinction time distribution. Specially we set *s* = 0.1 here. Panel (c) describes tumor extinction time distribution violin plots for parameter *d*, the natural death rate of effector cells. In the left half part (green color), effector cell and tumor cell binding rate parameter *γ* equals 0.02 and in the right half part (grey color), *γ* = 0.12. The inset shows the left two distributions in more detail. In panel (d), effector cell and tumor cell binding rate parameter *γ* is varied. The inset shows the right four distributions. In panel (e), the tumor killing rate *α*_1_ varies. For panel (f), the effector cell exhaustion rate *α*_2_ is varied. The inset zooms into the three distributions on the left. We fix other parameters *γ* = 0.12 and *α*_1_ = 0.5.

In contrast, the larger the tumor cell reproduction rate, effector cell death rate, or effector cell inactivation rate, the slower the tumor cell goes extinct. This also matches our intuition. We can also observe from Figure 5 that the increase of tumor reproduction rate and effector cell inactivation rate only slightly benefit the tumor survival by looking at the median extinction time.

We observe that some parameters influence extinction time distribution differently in the model without a conjugate compartment. The effector cell inactivation rate does not play an obvious role in tumor extinction time distribution. An increase of the two parameters *r* and *d* extends the tumor extinction time distribution, although the majority of tumor trajectories vanish at the beginning. Some parameters have opposite influence, parameters *s* and *β*_1_, representing effector cell reproduction rate and tumor cell killing rate; their increase leads to the shrink of the tumor cure time distribution. However, there are a number of tumor trajectories in the model without conjugate that take more time to be eliminated. Among the 1000 simulation runs, in Figure 6a, under *s* = 0.05 and *d* = 0.1 setting, four of the tumor trajectories fail to get extinction. Also, in Figure 6b, *r* = 0.3, *d* = 0.1 case, one run does not lead to tumor eradication.

**Figure. 6:**
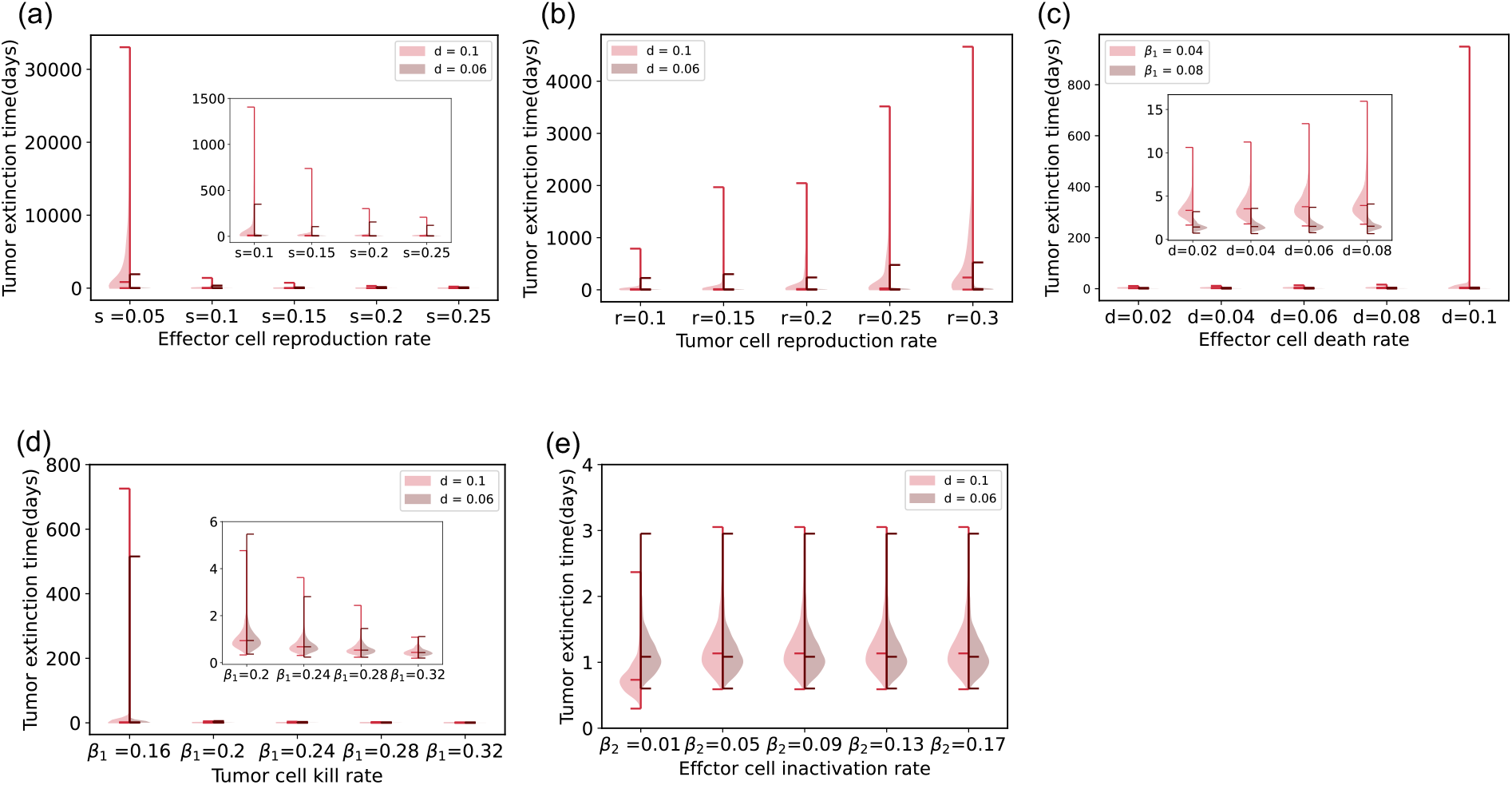
Stochastic simulation results - violin plots of tumor extinction time distribution for model without conjugate. The default parameters are *s* = 0.05, *r* = 0.15. In the left half part (pink), the effector cell death rate parameter *d* is set to be 0.1 and for the right half (brown), *d* = 0.06. The initial conditions are effector T cells *E* = 50, target tumor cells *T* = 100. Panel (a) displays tumor extinction time (in days) distribution violin plots for parameter *s*, the birth rate of effector cells. The inset zooms into the four distributions on the right. We set the other parameters *β*_1_ = 0.0175 and *β*_2_ = 0.0025. Panel (b) displays tumor cell reproduction rate *r* variability and extinction time distribution. Other parameters are fixed to *β*_1_ = 0.0178, *β*_2_ = 0.0022. Panel (c) describes tumor extinction time distribution violin plots for parameter *d*, the natural death rate of effector cells. In the left half part (pink rose color), effector cell and tumor cell binding rate parameter *β*_1_ = 0.04 and in the right half part (brown color), *β*_1_ = 0.08. The inset shows the left four distributions in more detail. We fix *β*_2_ = 0.005. In panel (d), tumor killing rate *β*_1_ varies. In the inset, we zoom into the four distributions on the right. Specially *β*_2_ = 0.05. In panel (e), the effector cell inactivation rate *β*_2_ varies. We fix *β*_1_ = 0.16.

## IV. DISCUSSION

The formation of cellular conjugates can crucially impact cancer-immune interactions on the cellular level. In this study, we investigated the importance of modeling the transition stage in tumor and effector cell interaction by comparing two models with and without explicitly considering the dynamics of a conjugate compartment. We characterized differences in the stability of the tumor and effector co-existence between these approaches. When conjugate dynamics are not explicitly considered but absorbed into the rates of killing and exhaustion, the tumor may reach equilibrium in stable coexistence with effector cells. However, this co-existence state can be altered if we explicitly consider the rate-adjusted equivalent model with conjugate compartment dynamics: the co-existence state can be unstable when the effector cell exhaustion rate is low, and the tumor-killing rate is also sufficiently low. For sufficiently large effector exhaustion rates, the tumor-effector co-existence state is stable for any tumor-killing rate.

In the model with a conjugate compartment, it takes more time to reach the stable tumor-effector cell coexistence state. The tumor and effector cell binding rate does not influence the damping rate of the oscillating ODE trajectories toward equilibrium, thus not influencing the time to reach the coexistence equilibrium state. However, the higher the tumor reproduction rate, the longer it takes to reach possible coexistence. Increasing any of the other four parameters (i.e., effector cell reproduction rate, effector cell death rate, tumor cell killing rate, and effector cell inactivation rate) reduces the time to converge to steady state. For the stochastic dynamics in the case of small population sizes, we found that parameters such as tumor effector cell binding rate, which did not affect the stability of deterministic co-existence, led to faster tumor extinction as it increased. Compared to the model with conjugate, parameters in the approach without conjugate show a more pronounced influence on the extinction time distribution.

Specially, we assumed the reproduction term of the effector cell population positively correlates to the tumor cell density. This results from effector cells only recruiting by antigen-presenting cells’ simulation which conveys information about tumor cell existence. This reproduction term makes our model different from the classical predator-prey model.

Our work suggests that the dynamics of an immune cell-tumor cell conjugate compartment can be crucial for the empirical understanding of possible tumor-control equilibria. The expected steady-state and stability may drastically change if this compartment is not explicitly considered. A possible co-existence state is unstable as its behaviors largely depend on the explicit parameter values in the model with a conjugate compartment. Thus, including a conjugate compartment, tumor populations can escape if the tumor-killing rate is sufficiently low. If tumor-the dynamics of the transient conjugate compartment are somewhat ignored because immune conjugates are assumed to be in a quasi-steady state [29], the rate of tumor escape might be more challenging to predict. The pioneering work by Kuznetsov et al. [29] focused on the quasi-steady-state model of the cytotoxic T lymphocyte response to the growth of tumor, assuming the conjugate dynamic is constant. The conjugate formation process before the effector T cell kills target tumor cells can be regarded as a handling time term for predator and prey models. The handling time here also defines how effector killing is reduced due to effector cell binding with a target tumor cell. Modeling the tumor-immune conjugation as a separate compartment allows for more flexibility regarding the possible outcomes of cancer-immune predation, such as tumor escape or tumor control.

Understanding the critical conjugation process in tumor–immune interactions will benefit the development of immunotherapies for predicting long-term outcomes such as tumor control or eradication. Our mathematical model provides a framework to describe the dynamics of small tumor populations emerging during early cancer evolution or after treatment, resulting in either extinction or progression. The work presented here takes a step toward demonstrating the importance of the conjugate compartment in defining tumor-effector interactions. Models of early-stage onco-immunological interactions taking transitionary compartments into account, may better capture the dynamics of tumor progression. For future modeling geared toward clinical data, considering the conjugate compartment may lead to improved insights in predicting tumor fates.

## DATA AVAILABILITY

The simulation code for this project is publicly available, https://github.com/paltrock/conjugates.

## ACKNOWLEDGMENTS

The authors acknowledge funding from the Max Planck Society, the DFG Research Training Group TransEvo (GRK 2501, Project number 400993799), and are grateful for fruitful discussions with members of the Theoretical Biology Department at the Max Planck Institute for Evolutionary Biology.

## AUTHOR CONTRIBUTIONS

All authors contributed to the study’s conceptualization, methods, formal analyses, writing, and manuscript editing. QY executed the mathematical and computational work, developed the mathematical model, and performed statistical analyses. QY and PMA wrote the manuscript with input from AT.

## Appendix A: Stability of steady states

### 1. Model without conjugate

The model without conjugate, Eq. (3) has two steady states. Here we prove that the non-trivial one is stable. The Jacobian is given by 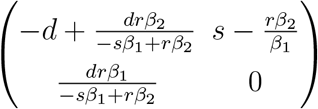. The eigenvalues *λ*_1_ and *λ*_2_ are

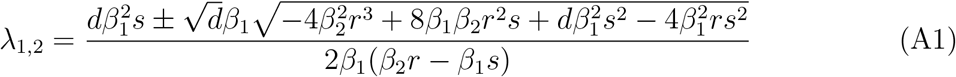

We show that the real part of both eigenvalues is always negative. The fixed point only exists if *β*_2_*r < β*_1_*s*, cf. II. Therefore, the denominator is always negative. Thus, we have to show that the real part of the numerator of *λ*_1,2_ is always greater or equal to 0.

There are two cases:

i. If the argument of the square root, 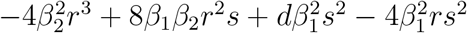, is negative, the real part of the two Eigenvalues is given by

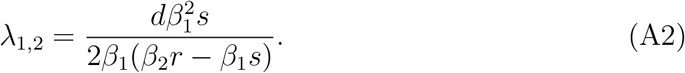

As the denominator is negative, the real part is negative.
ii. If the argument of the square root, 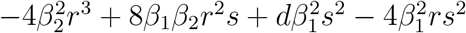, is positive, we need to show that

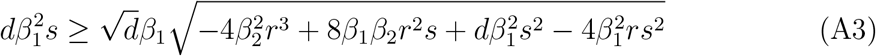

We divide by 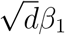

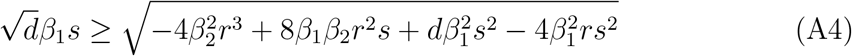

and square both sides

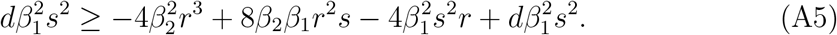

We remove the term 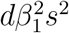 to arrive at

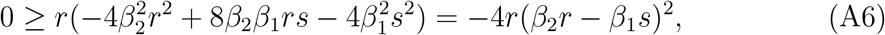

which is always fulfilled. Thus, the numerator is always positive and thus the Eigen-values have always negative real part.

### 2. Model with conjugate

For the model with conjugate, we numerically solve the ordinary differential equations and plot the bifurcation diagram for tumor and effector cell populations and parameter tumor cell killing rate *α*_1_, shown in Figure 7. We vary *α*_1_ from 0.2 to 1, to see the populations and the stability change.

**Figure. 7:**
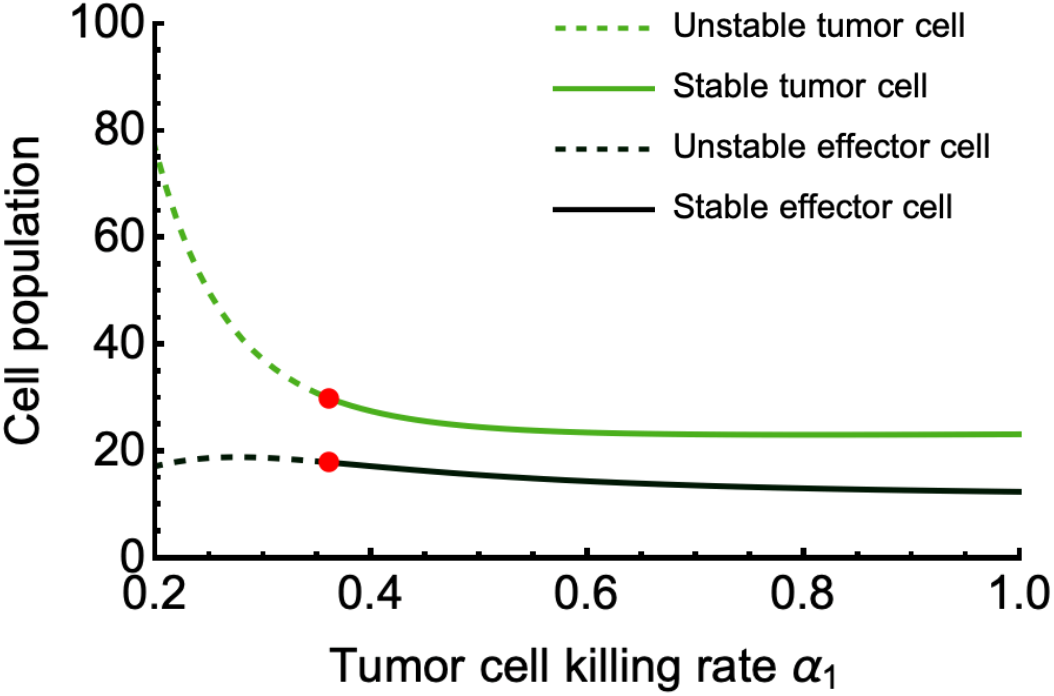
Bifurcation plot. The tumor cell population is green, and effector cell population is dark green. Dashed lines are unstable fixed points and lines are stable fixed points. Red points indicate the bifurcation points (parameters *s* = 0.05, *r* = 0.15, *d* = 0.1, *γ* = 0.01, and *α*_2_ = 0.01).

